# Social-vocal brain networks in a non-human primate

**DOI:** 10.1101/2021.12.01.470701

**Authors:** Daniel Y Takahashi, Ahmed El Hady, Yisi S Zhang, Diana A Liao, Gabriel Montaldo, Alan Urban, Asif A Ghazanfar

**Affiliations:** Princeton Neuroscience Institute, Princeton University; Princeton, USA; Department of Psychology, Princeton University; Princeton, USA; Department of Ecology and Evolutionary Biology, Princeton University; Princeton, USA; Brain Institute, Federal University of Rio Grande do Norte; Natal, Brazil; Department of Collective Behavior, Max Planck Institute of Animal Behavior; Konstanz, Germany; Center for Advanced Study of Collective Behavior, University of Konstanz; Konstanz, Germany; euro-Electronics Research Flanders; Leuven, Belgium; VIB; Leuven, Belgium; Imec; Leuven, Belgium; Department of Neuroscience, Faculty of Medicine, KU Leuven; Leuven, Belgium

## Abstract

During social interactions, individuals influence each other to coordinate their actions. Vocal communication is an exceptionally efficient way to exert such influence. Where and how social interactions are dynamically modulated in the brain is unknown. We used functional ultrasound imaging in marmoset monkeys – a highly vocal species - to investigate the dynamics of medial social brain areas in vocal perception, production, and audio-vocal interaction. We found that the activity of a distributed network of subcortical and cortical regions distinguishes calls associated with different social contexts. This same brain network showed different dynamics during externally and internally driven vocalizations. These findings suggest the existence of a social-vocal brain network in medial cortical and subcortical areas that is fundamental in social communication.

**One Sentence Summary:** A network of medial subcortical and cortical brain areas gate social communication in primates.

---

Vocal behavior is special relative to other forms of communication in that it can be used to rapidly influence the behavior of others at a distance or even out-of-sight from other group members. In some cases, it strengthens social bonds by maintaining contact (*1, 2*); in other cases, it warns of danger (whether intentionally or not) (*3*). Variations embedded in the acoustics of vocalizations allow listeners to distinguish between different types of utterances (e.g., contact or alarm vocalizations) and indexical cues (e.g., identity, body size and/or age) (*4*). These acoustic cues help listening individuals make decisions; for example, whether or not to reply with a vocalization, to keep silent, to move towards the source or away.

Naturally, investigations into the mechanisms of vocal behavior focus largely on how, in a listener’s brain, signals from the ear are processed by specialized circuits in the auditory system (*5, 6*), sent via direct connections to the frontal cortex (*7, 8*) whereby their activities modulate behaviors (*9-11*). However, in parallel to this cortical transformation of auditory signal to motor act in vocal communication are subcortical neural processes that are largely unknown and likely shared among many types of adaptive behaviors regardless of modality. Across all vertebrates, a group of six highly conserved midbrain and brainstem areas is critical to produce different social behaviors (e.g., mating, aggression, parental care). Dubbed the “social behavioral network” (SBN) it consists of the anterior hypothalamus (AH), pre-optic area (POA), ventromedial hypothalamus (VMH), lateral septum (LS), periaqueductal gray (PAG), and extended amygdala (ExtAm) (*12-14*) (fig. S1A). The SBN is thought to functionally assemble in different ways to produce different adaptive social behaviors. Although this parallel has not been made explicitly in the literature, much of the SBN overlaps with subcortical regions involved in primate vocal production (*15, 16*). Given their dense connectivity with frontal cortical regions (*17*), understanding how the SBN interacts with frontal cortical areas is a necessary step to understanding the evolution of the primate social communication (*17, 18*).

We used functional ultrasound imaging (fUS) in the brains of marmoset monkeys to investigate the role(s) of the SBN in vocal perception, production, and audio-vocal interaction. In addition, we related SBN activity to a key frontal cortical region involved in vocal production: the anterior cingulate cortex (ACC) (fig. S1A) (*19*). Our goal was to determine how the SBN is involved in vocal communication and how networks were dynamically and differentially assembled as a function of different call types during vocal perception and production. Marmoset monkeys are particularly suitable for this investigation because they are highly social species, producing a rich repertoire of vocalizations whose production is a function of social context (*20, 21*). We used fUS--which measures changes in cerebral blood volume (a proxy for neural activity; (*22, 23*))--because it allowed for a wide coverage (16×20mm^2^) of the medial brain region, capturing activity in most of the SBN and medial frontal cortical areas at high spatial resolution (∼125×200×400 µm^3^) and with a temporal resolution fast enough (2 Hz) to image brain dynamics in vocalizing and alert marmosets (fig. S1B).

Studies investigating the auditory responses to vocalizations in regions associated with mammalian vocal production are few though songbird neurobiology suggests that certain specialized brain regions may have auditory-vocalization function, e.g., HVC (*24*). Hence, we first asked: Do the SBN and ACC respond auditorily to vocalizations? Using a block design, we played back five different vocalizations and one amplitude-modulated noise signal to marmosets (n=5) while imaging their midline brain regions. Two vocalizations were long-distance contact calls (phee and twitter), two were short-distance contact calls (trillphee and trill), and one was an alarm call (*25*); the noise signal was white noise amplitude-modulated to match phee calls (Fig. 1A). Each call block lasted for 18 s with 25 to 50 s intervals between the blocks (Fig. 1B). The total amount of acoustic energy was matched between blocks. To localize specific brain areas, we registered the fUS images to a marmoset brain atlas (Fig. 1C). Figure 1D shows an example of fUS signal acquired from ACC during a single session for one marmoset. Figure 1E shows some examples of average CBV responses to vocalizations, with large responses after the call block onset (red lines indicate significant activity compared to the CBV activity before the playback onset). We plotted the areas in which the peak activity was significantly different from the baseline (p <0.05, Bonferroni corrected) (Fig. 1F): large portions of subcortical areas and the medial prefrontal cortex were significantly activated. The ACC and regions within the SBN exhibited the strongest brain responses to vocalizations. Limbic thalamus (LT) (see Fig S1A) also showed a significant response consistent with its role in connecting SBN and ACC (*26*).

**Fig. 1:**
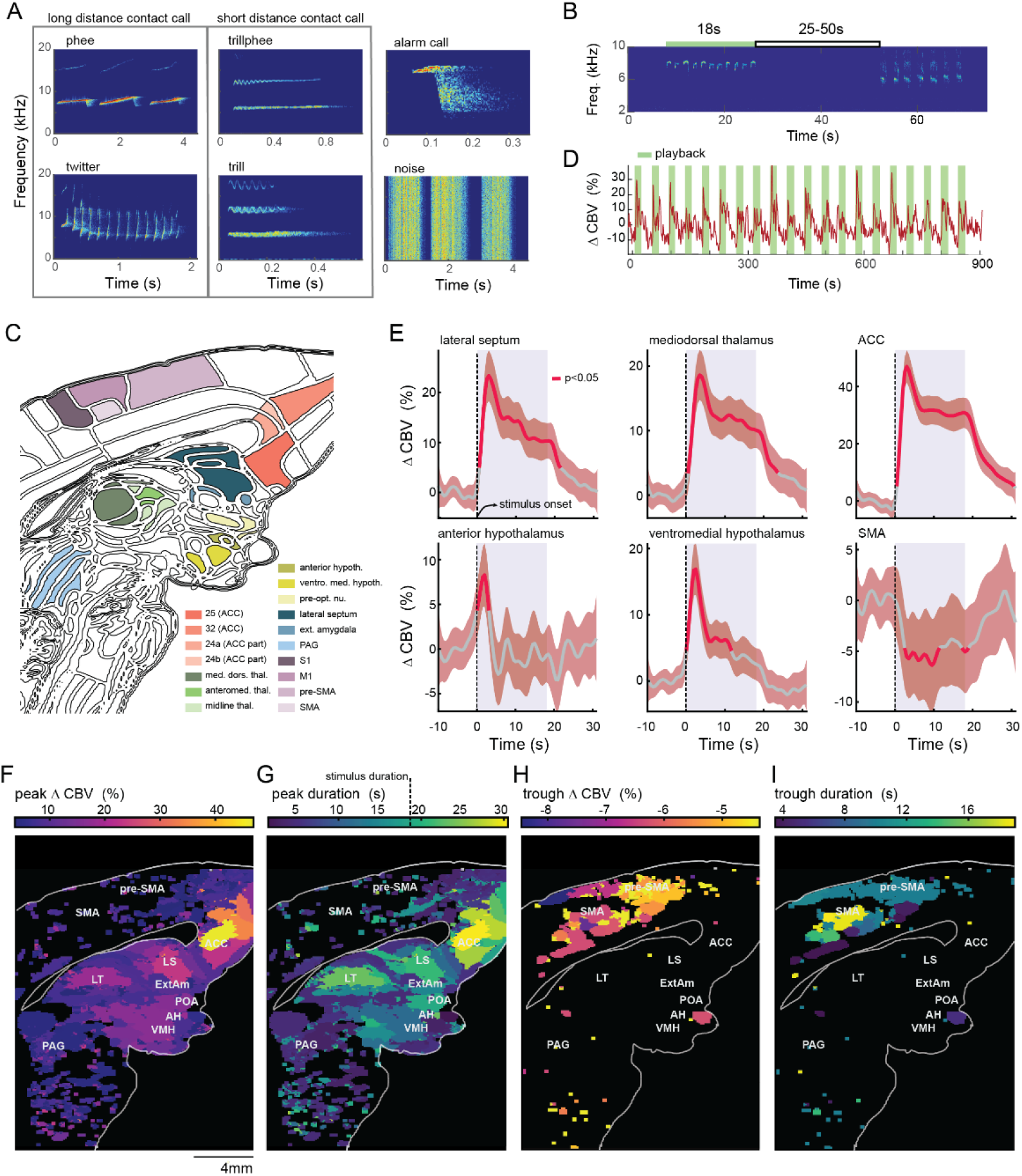
Brain dynamics during vocal perception: **(A)** Spectrogram of the different call types used in playback. **(B)** Spectrogram of two example block stimuli. **(C)** Annotated atlas registered to a reference fUS. **(D)** An example of the CBV response in the ACC. Green stripes represent the time during which the stimuli were played. X-axis: percentage change of CBV. **(E)** Example brain regions that showed significant response while the marmoset was listening to the playbacks (dotted line and shaded blue area represent the onset and duration of the stimulus, respectively). Solid red lines indicate CBV activity that was significantly different (p <0.05 Bonferroni corrected) from the baseline activity (−10 to 0.5 s before the stimulus onset. Shaded red indicates 95% confidence interval. The CBV response had peaks and troughs. **(F)** Activity map of medial brain regions showing areas that had the significant maximum peak in CBV in response to playback (p<0.05 FDR corrected). Colorbar indicates the percentage difference between baseline and peak response. **(G)** Duration map of medial brain regions showing the duration of the brain response. Colorbar indicates the duration of the significant peaks in (F). The dotted line in the colorbar indicates the stimulus duration (18 s). **(H)** Activity map of medial brain regions showing areas that had significant trough (largest decrease) in CBV in response to playback stimuli (p < 0.05 FDR corrected). **(I)** Duration map of medial brain regions showing the duration of the brain response (significant CBV troughs).

Figure 1E shows that some areas exhibit sustained activity (e.g., ACC), other areas have short-lived activities (e.g., anterior hypothalamus), and still others have decreased responses (e.g., SMA). Sustained brain activation are thought to modulate behavior even after the stimulus is no longer present, whereas short-lived activations are thought to be related to perceptual processes (*27*). Additionally, areas with decreased brain activity shows that these areas are deactivated, possibly gating sensory-motor information during acoustic perception (*28*). Several SBN areas (e.g., LS, POA) and ACC showed longer sustained activity (Fig. 1G). Medial sensorimotor cortex (mSMC) had significant CBV decrease in response to the playback, especially in the supplementary motor area (SMA) and pre-supplementary motor area (pre-SMA) (Fig. 1H-I). These areas exhibited CBV dynamics that were negatively correlated with the SBN and ACC activity (fig. S2). These findings indicate that SBN and ACC process sensory information and mSMC suppresses motor activity during vocal perception.

A key prediction of SBN activity is that the network should respond differently in distinct social contexts (*12, 14*). We could test this prediction by comparing the brain response to different call types. If brain activity is related to the context in which each calls are produced, we expected that long-distance contact calls (phee and twitter calls) would exhibit similar CBV responses that are different from the responses to both short-distance contact calls (trillphee and trill calls) and alarm calls. Conversely, if the brain responses are related straightforwardly to simpler acoustic characteristics of the calls (like a primary auditory region), we expected that the response to phee calls and trillphees should be more similar compared to other calls, because their acoustic characteristics are more similar (fig. S3A-B). We also used responses to the amplitude-modulated white noise as a control for phee calls (same duration and amplitude modulation, but different frequency content) to verify that amplitude modulation and call duration are not the principal factors for the brain response.

Figure 2A shows that CBV response to phee and twitter are statistically indistinguishable for all medial brain areas (p > 0.05 FDR corrected). In contrast, responses to trillphee, trill, and alarm calls exhibit significant differences from the phee, especially in the SBN and ACC (Fig. 2B-D, p< 0.05 FDR corrected, fig. S4). Amplitude and duration-matched noise stimuli showed the largest differences among stimuli, having no natural communicative content (Figs. 2E, p < 0.05 FDR corrected). To further quantify the degree of similarity between brain responses to different types of calls, we calculated the hierarchical clustering of CBV activity across the entire medial brain region. If the medial brain regions integrate the social context associated with vocalizations, we expected that responses to long-distance contact calls would be grouped together. In contrast, responses for short-distance contact calls should be grouped in a different cluster, as was indeed the case (Fig. 2F).

**Fig. 2:**
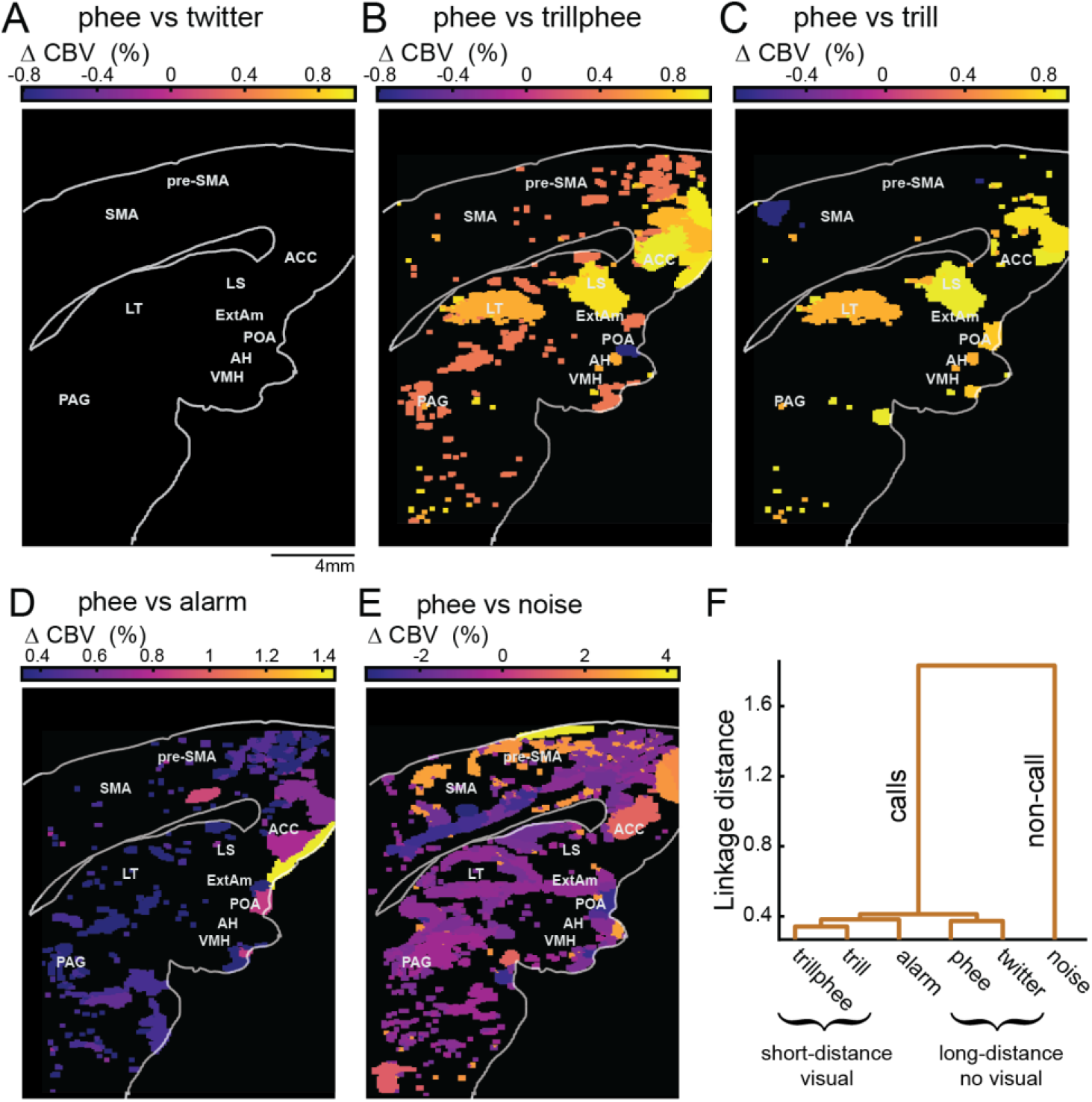
Comparing medial brain dynamics for different call types during vocal perception. Maps showing medial brain regions with significantly different activity (p < 0.05 FDR corrected) between **(A)** phee and twitter, **(B)** phee and trillphee, **(C)** phee and trill, **(D)** phee and alarm call, and **(E)** phee and noise. Colorbars indicate the difference between CBV response to phee and the respective calls. Negative values indicate that that CBV was larger for phee. Regions with black color indicate areas that were not statistically significant. (**F)** Hierarchical clustering of all activity maps in response to different call types.

Given that vocal communication is a sensory-motor interaction (*29*), to fully understand the role of SBN during vocal communication we also need to study its dynamics during vocal production. Many physiological and lesion studies have implicated parts of the SBN and medial frontal cortex as important for vocal production (*30-32*). However, how those different areas interact during vocal production -- perhaps differentially assembling for different vocalizations -- is unknown. We used fUS to image CBV changes in the medial brain region in marmosets while they produced different vocalizations (phees, trills and alarm calls; n=3 of the 5 subjects used for the perception experiment).

Figure 3A shows an example of a single contact call production and the corresponding CBV change in ACC. More generally, the average dynamics of medial brain activity during vocal production varies a lot among areas (Fig. 3B). To better understand these dynamics, we plotted the significant largest peak CBV activities (p < 0.05, Bonferroni corrected, Fig. 3C). Large parts of the medial region showed significant activation during vocal production. Brain areas related to vocal initiation exhibit the strongest increase in CBV activity before the onset of spontaneous vocal production (defined behaviorally as a vocalization produced following at least a 12s period of no vocal production from the subject or a nearby conspecific (*29*)). On the other hand, areas related to sensory-motor modulation of ongoing vocalization have the largest peak activity during the spontaneous vocalization. By calculating the timing of the largest CBV peak with respect to the onset, SBN areas showed activation before the onset, whereas mSMC showed a CBV peak immediately after the onset (Fig. 3D). The ACC showed a delayed CBV response compared to the rest of the medial brain regions, indicating a different role during vocal production (Figs. 3B,D). These findings suggests that (1) SBN areas are related to preparation and initiation of spontaneous vocalization, (2) mSMC is related to sensory-motor modulation of ongoing vocalization, (3) ACC is perhaps involved in monitoring the outcome of vocalization(*33*).

**Fig. 3:**
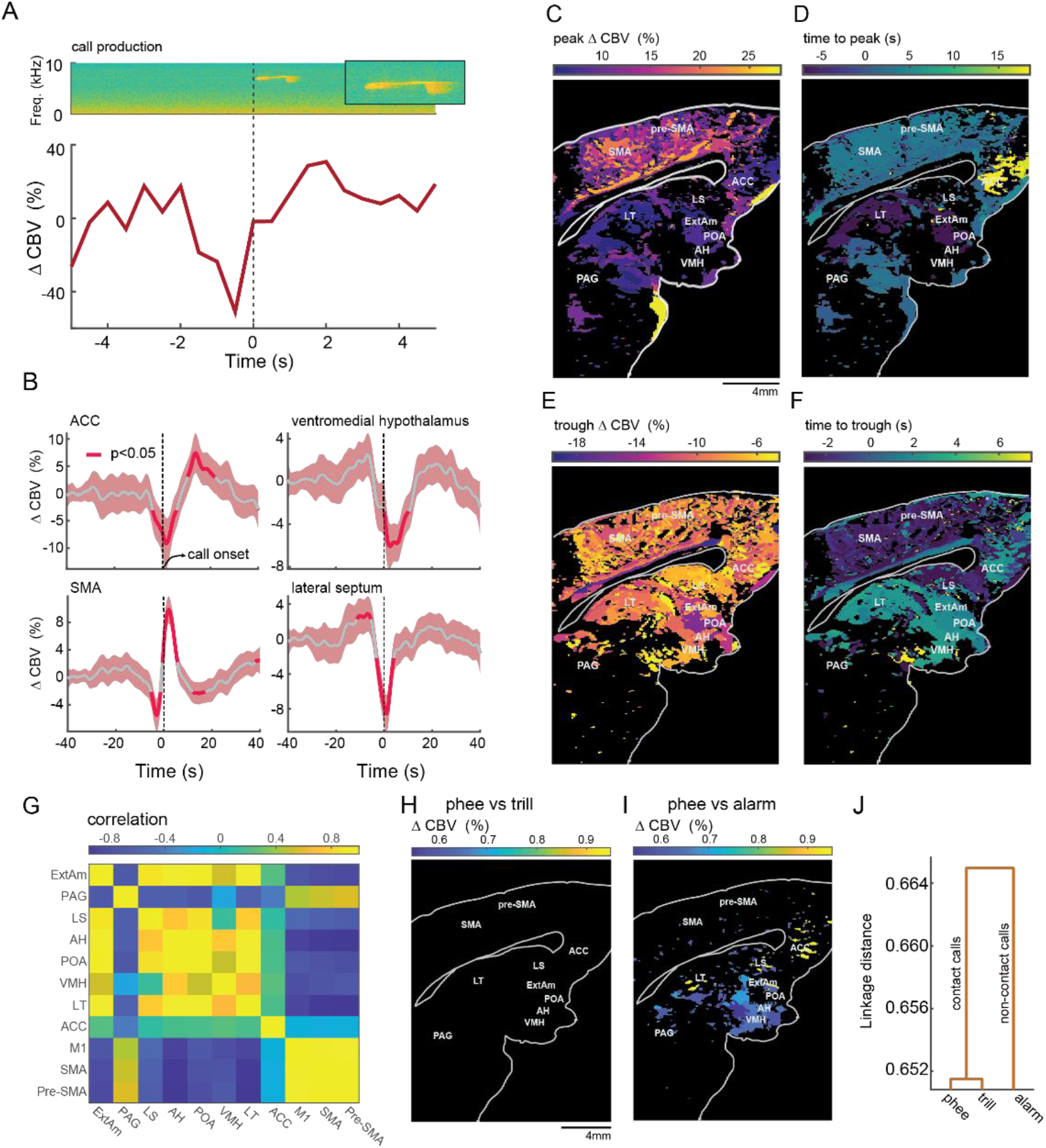
Medial brain regions activity during vocal production. **(A)** Example of CBV activity during a single contact-call production in ACC. Top: spectrogram of the call. Bottom: corresponding CBV. **(B)** A subset of brain areas shows changes in the CBV during vocalization. Average CBV activity for all call types. Solid red lines indicate CBV activity that was significantly different (p<0.05 Bonferroni corrected) from the baseline activity (−40 to −30 s before the call onset). Shaded red indicates 95% confidence interval. **(C)** Activity map showing the brain regions that have significant peak in the CBV (p < 0.05 Bonferroni corrected). Colorbar shows the difference between baseline and peak. **(D)** Delay map showing the brain regions and the time to peak in the areas shown in (C). Colorbar shows the delay to the maximum peak. **(E)** Activity map showing brain regions that have significant decrease in the CBV (p < 0.05 Bonferroni corrected). **(F)** Delay map showing the time to the trough of the CBV of the areas in (E). Colorbar shows the delay to the lowest trough. **(G)** Correlation between SBN, ACC, SMA, pre-SMA, and, M1 activity. **(H) and (I)** Maps showing medial brain areas that have significant differences (p < 0.05 FDR corrected) in the CBV activity between **phee** and trill calls production and phee and alarm calls production, respectively. Colobar shows the difference between CBV activity during phee and the respective calls. **(J)** Hierarchical clustering of all brain activities in response to phee, trill, and alarm calls showing clusters of contact calls versus non-contact calls.

Areas that exhibit an increase in CBV activity can also show a decrease in CBV activity at different time points relative to the call onset (Figure 3B). A natural question is how the increase and decrease in CBV activities are coordinated among different brain areas. One possibility is that SBN and ACC show CBV dynamics that alternates with the dynamics of mSMC, similar to the CBV dynamics during vocal perception. To test this hypothesis, we plotted the areas with activity troughs significantly different from the baseline (p < 0.05, Bonferroni corrected, Fig. 3E). Again, the SBN, ACC, and mSMC showed significant suppression. When we computed the timing of the suppression with respect to the onset of calls, mSMC showed it before the onset of call and, in general, SBN and ACC exhibit suppression after the call onset (Fig. 3F). This suggests a negative correlation between SBN+ACC and mSMC activity. We confirmed this by calculating the correlation matrix between SBN, ACC and medial cortical motor areas (Fig. 3G, fig. S5). The only exception was PAG which was correlated positively with cortical motor areas and negatively with SBN, consistent with its role as a motor area during vocalization (*31*).

Next, we wanted to know whether the SBN and ACC differentiated between different spontaneously produced vocalizations. If SBN and ACC activity is related to social context in which calls are produced, we predict that the SBN and ACC activity for vocal production is similar between contact call types (phees and trills) versus alarm calls. In support of this, a comparison of CBV response trajectories between contact calls (phees versus trills) did not exhibit a significant difference (Fig. 3H), but phee versus alarm call production showed significant differences in SBN and ACC regions (Fig. 3I). Moreover, when hierarchically clustered, brain activities during contact calls production were grouped together whereas brain activities for alarm calls were clustered separately supporting the notion that the medial brain regions are social context-sensitive (Fig. 3J).

A social context is not defined only by the usage of different types of calls, but also by the history of vocal interaction. A spontaneous vocalization can be distinguished from a vocalization produced in response to another’s vocalization (externally-driven; e.g., vocal turn-taking; (*29, 34*) or one that was immediately preceded by the vocalization from the same individual (internally-driven; e.g., a vocal sequence (*29*)). If the medial brain processes social context encompassing both past vocal history and vocalization types, we predicted that there would be a difference in the brain dynamics during vocal production depending on whether a call is spontaneous, in response to a conspecific’s call or preceded by a vocalization from the same individual. We also hypothesized that the effect of call history dependence would be more salient for contact phees and trills, which are often produced during vocal interaction (*20*), than for alarm calls. Figure 4A shows that regions of the SBN and ACC areas are more activated when produced in response than when produced spontaneously (p < 0.05 FDR corrected). This result is consistent with the hypothesis that SBN+ACC network functions to process externally-driven social contexts adaptively. Figure 4B shows that the mSMC is more strongly activated during sequences of calls when compared to spontaneous calls (p < 0.05, FDR corrected), indicating that mSMC is related to internally driven vocalizations. We observed significantly less areas with history dependence for alarm calls (figs. S6A-B).

**Figure 4:**
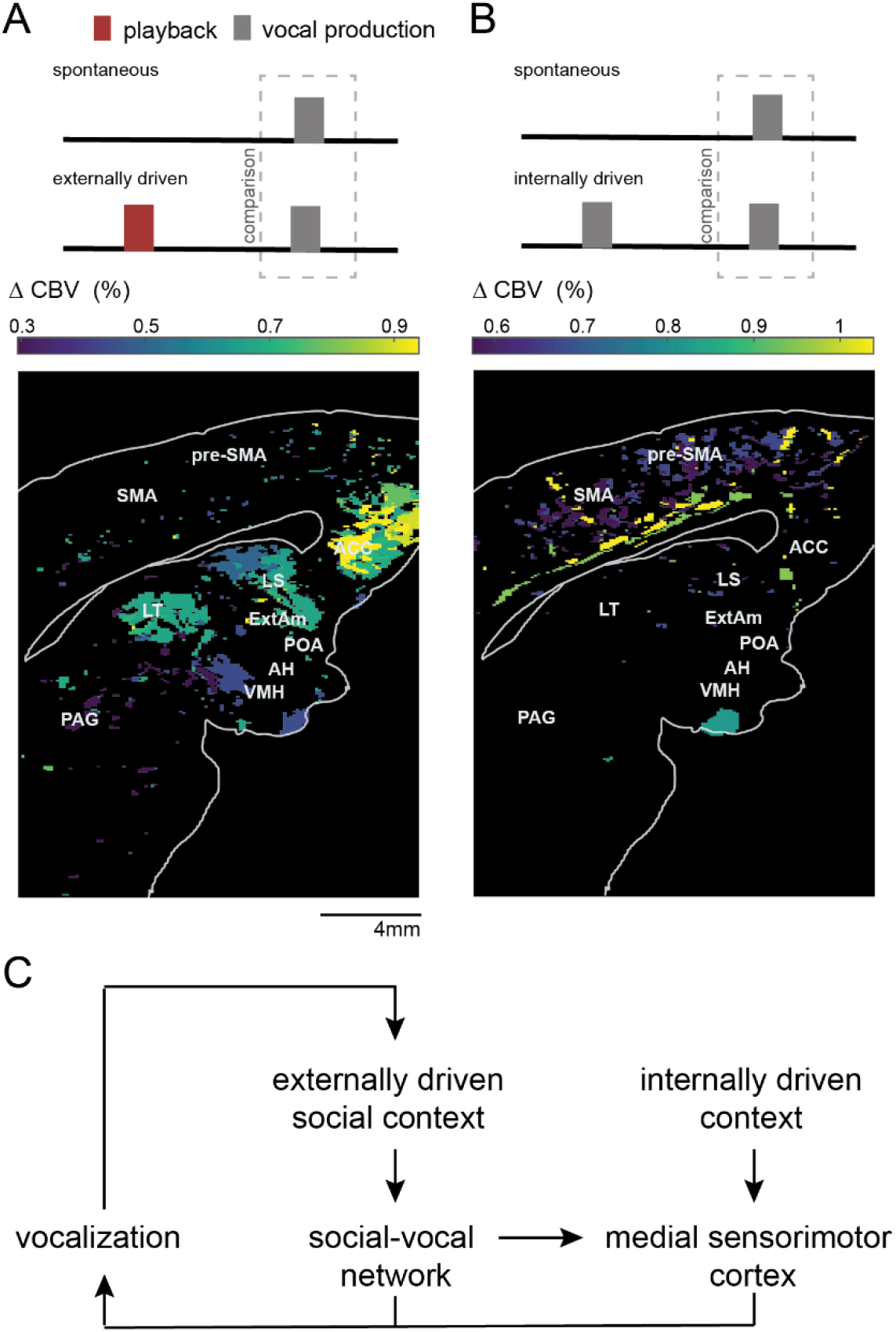
History dependence in the medial brain regions during audio-vocal interaction. **(A)** CBV activity differences between spontaneous and response contact calls. The brain map shows significantly different areas between both conditions (p < 0.05 FDR corrected). Colorbar shows the percentage difference between the CBV activity during production of (externally driven) response and spontaneous contact calls. Positive values indicate that the activity was stronger for the response to playback calls. **(B)** CBV activity difference between spontaneous and sequence contact calls. The brain map shows significantly different areas between both conditions (p < 0.05 FDR corrected). Colorbar shows the percentage difference between the CBV activity during (internally driven) sequence and spontaneous call production. Positive values indicate that the activity was stronger for sequence contact calls. **(C)** Proposed model of the effect of externally driven social context on social-vocal network (SBN, ACC, LT) and of internally driven context on mSMC.

During a conversation, an individual initially produces vocalizations adapted for the social context. Then another individual listens to the vocalizations until it finishes before responding. While listening, the same individual integrates the social contexts to produce the response that can be a single or a sequence of vocalizations depending on the internal state. The repetition of this cycle creates complexity in the vocal communication going beyond what can be accomplished with an isolated utterance (*35*). Our results suggest a neural mechanism for such a communicative cycle (Fig. 4C). During vocal perception, SBN and ACC encode social context. At the same time, mSMC is suppressed so that other sensory-motor activities like body movements are avoided, making the marmoset listen to the calls. Once the subject listened to the calls, SBN and ACC activities will influence the response call production. To produce a call sequence, the mSMC is activated, similar to the execution of other sequential behaviors (*36*).

Previous studies have shown that SBN is a key area modulating social behaviors in fishes and tetrapods (*12, 14*). Interestingly, in primates, including humans, most research on the social brain concentrated on the role of cortex in the social cognition, partly due to methodological choices and partly due to the salience of the cortex in primates (*17, 37, 38*). Consequently, there is a lack of studies of SBN in primates during social cognition, despite the hypothesis that it should be universally relevant for social behavior in tetrapods. On the other hand, when SBN areas are studied in primates, they are often associated with “simpler” roles like initiation or production of species-typical behaviors (as pointed out by (*17, 39*)). For instance, in vocal communication, brain areas that constitute SBN are considered part of primary vocal motor network related to the initiation and production of “emotional” vocalizations, contrasting with lateral frontal cortical areas which are associated with cognitive control of vocalization (*15, 40*). Naturally, this led to the hypothesis that the evolutionary change in primate social communication (especially in humans) is driven mainly by cortical areas (*41*). We show that SBN in marmoset monkeys has a role beyond simple production of vocalizations, being related to social perception, vocal modulation, and historicity in vocal communication, all of which generate flexibility in social interactions. This conclusion is also bolstered by the fact that communication in non-mammalian “cortex-less” vertebrates also show a degree of sophistication not expected (*42*). Together, these results support the hypothesis that the tinkering of SBN and its connections with frontal cortical areas like ACC were a key step in the evolution of primate communication, including human speech (*18*).

## Author Contributions

Conceptualization: DYT, AEH, AAG

Experimental design: DYT, AEH, AAG

Funding acquisition: AAG

fUS probe design: GM, AU

Data collection: DYT, AEH, YSZ, DAL

Data analysis: DYT, AEH

Image registration: YSZ

Writing – original draft: DYT, AEH, AAG

Writing – review & editing: DYT, AEH, AAG

## Supplementary Materials

### Material and Methods

#### Subjects

The subjects used in this study included five adult common marmosets (*Callithrix jacchus*) housed at Princeton University. The marmosets were three males and two females. Animals were fed once daily with standard commercial chow supplemented with fresh fruits and vegetables. The animals had ad libitum access to water. The colony room was maintained at a temperature of approximately 27C and 50%– 60% relative humidity, with a 12 hr light:12 hr dark cycle. Before experiments were conducted, all animals were familiarized with the testing room and imaging equipment. All experimental sessions were performed in compliance with the guidelines of the Princeton University Institutional Animal Care and Use Committee.

### Surgery

Initial monitoring included temperature, pulse, respiration, and SPO2. Blood was collected for pre-operative glucose measurement. Dexamethasone 1 mg/kg IM and Baytril 5mg/kg IM were administered pre-operatively. The animal is induced with alfaxalone 10 mg/kg IM and intubated. The animal is carefully placed in a marmoset-specific stereotaxic device, and the exposed skin is prepared. All the following procedures were executed in sterile conditions. The skull overlying the brain region of interest was exposed by making an incision along the top of the head through the skin. The tissue was reflected, and the periosteum removed until the skull was exposed. Sterilized miniature titanium screws were inserted into the bone at various positions to serve as anchors to hold the head plate to the skull covering the exposed area’s border. The head plate is a machined piece of flat metal with a rectangular hole in the center that allows the fUS probe to be attached and aligned during imaging experiments. The head plate is attached to the skull and screws with dental cement (C&B Metabond® Quick Adhesive Cement System, Parkell). A head post was fixed to secure the hardware. A cranial window of ∼ 8 mm X 16 mm was created with a piezoelectric drill that does not damage soft tissues. To protect the craniotomy when the animal was not being imaged, the cranial window was sealed using silicone gel (Kwik–cast; World Precision Instruments). A stainless-steel cover was secured headplate to cover the headplate and headshield.

### Functional ultrasound imaging (fUS)

To measure the hemodynamic change of the brain areas in alert and vocalizing marmosets, we used functional ultrasound imaging (fUS) (*22, 42*). The hemodynamic change measured by fUS strongly correlates with the cerebral volume change (CBV) change of the arterioles and capillaries. It compares more closely to CBV-fMRI signal than BOLD-fMRI (*23*). CBV signals show a shorter onset time and time-to-peak than BOLD signals in marmosets (*43*). fUS signals correlate linearly with neural activity for various physiological regimes (*44, 45*). We used a custom ultrasound linear probe with a minimal footprint (20mm by 8mm) and light enough (15 g) for the animal to carry. The probe comprises 128 elements of 125µm pitch working at a central frequency of 12MHz, allowing a wide area coverage (20mm depth, 16mm width). The probe was connected to an ultrasound scanner (Vantage 128, Verasonics) controlled by an HPC workstation equipped with 4 GPUs (AUTC, fUSI-2, Estonia). The functional ultrasound image formation sequence was adapted from (*42*). The main parameters of the sequence to obtain a single functional ultrasound imaging image were: 200 compound images acquired at 500 Hz, each compound image obtained with 9 plane waves (−6° to 6° 1.5° steep). With these parameters, the fUS had a temporal resolution of 2Hz and a spatial resolution (point spread function) of 125*μ*m width, 130*μ*m depth, and 200 to 800 *μ*m thickness depending on the depth (200*μ*m at 12mm depth). We acquired fUS signal at the midline sagittal plane (0mm).

### Relation between fUS signal and cerebral blood volume

The Doppler intensity is proportional to the part of the CBV signal corresponding to red blood cells moving faster than ∼1 mm/s in the z-direction (*CBV*_*filter*_):

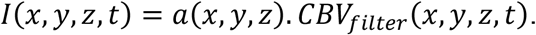

The constant of proportionality is different for each voxel and depends on several parameters, including emitted power, tissue attenuation, the geometry of the probe, etc. To remove such constant, we defined the hemodynamic signal measured by fUS as Δ*CBV* representing the variation of the Doppler intensity compared to the background signal:

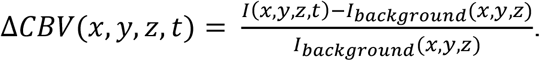

The background signal was the low-frequency component (< 0.01Hz) of the signal.

### Experimental protocol for fUS recording

Each animal was tested only once per day at around the same time of the day each time. First, the experimenter transfers the marmoset from its housing to the experimental room. The experimental room measured 3.2m x 5.5m with walls covered in sound attenuating foam. A table (0.75 m in height) was positioned at one of the corners of the room. The animal is put in a custom-designed partial restraint device, and the head is temporarily fixed using the head post. Afterward, all procedures were performed with sterile surgical gloves as well as autoclaved tools and materials. The head cover is then removed to expose the cranial window. The experimenter flushes the cranial window using sterile water and applies de-bubbled sterile ultrasound gel (Sterile Aquasonic 100 Ultrasound Gel) onto the recording surface. A custom-designed probe holder is fixed on top of the head plate. The ultrasound probe is then placed inside the probe holder. The experimenter releases the head post so that the animal has a wider range of movement. The experimenter took an initial functional ultrasound image to ensure no large movement artifacts and that selected the right plane. Marmosets were then recorded during the experiment for ∼1hour. After the recording, the experimenter cleans the recording surface with chlorhexidine 0.05% and sterile water. Then a new sterile headcover is secured. The marmoset is removed from the chair afterward and is returned to its housing.

### Acoustic recordings

A microphone (Sennheiser ME66) was positioned near the top of the animal at a 60cm distance from the top of the partial restraint device. This microphone was connected to a ZOOM H4n Handy Recorder, which worked as a digital amplifier. Audio signals were acquired at a sampling frequency of 96 kHz. fUSi signal and acoustic signals were synchronized using the TTL signal from the ultrasound machine, indicating image formation.

### Playback procedures

A speaker (JBL LSR305 5”) was located on the other side of the room at 4m from the animal. An opaque cloth occluder prevented the subjects from visualizing the speaker. We calibrated the sound level to be at 60dB at the animal head location. Ambient noise was below 30dB. The fUSi recording session always started with 10 minutes recording of baseline CBV activity without playback stimuli. For the acoustic stimuli we used 5 different call types (Phee, trillphee, trill, twitter, and alarm call) with one scrambled noise stimulus. These were presented using a block design with a playback block duration of 18s. The between block intervals ranged between 25 s to 50 s and the duration were randomly and uniformly chosen for each trial. Each block was composed of the same call type from the same subject. We recorded calls from nine marmosets with more than 20 calls for each type of call. Each subject only received playback calls of the animals other than self. We randomized the order of the type of call of the block and the calls chosen within each block. The total acoustic power within a block was normalized to be the same for every block. The interval between calls within the same block was one second.

### Vocalization data

Marmoset vocalization and fUS signals were simultaneously recorded while an animal vocalized spontaneously (*i*.*e*., no playback or self-produced vocalization within 12 s before the vocalization onset), as a response to playback stimuli (*i*.*e*., playback offset was within 12 s to 0.5 s before the vocalization onset), and as a sequence to another call produced by the subject (*i*.*e*., subject’s previous call offset was within 12 s to 0.5 s before the next vocalization onset). We detected call using Audacity spectrogram view. The onset and offset of a call were defined as the first and last time points with the power at the fundamental frequency above the background noise. Automated detection programs had a considerable false-negative rate and were not reliable, especially for soft calls. Onset and offset of vocalization were defined as the first and last time points at which the spectrogram of the call was visible.

### Trials exclusion criteria

To detect movement artifacts, we calculated the average fUS signal for each frame. The trial was excluded if the average fUS signal for a frame during a playback or vocalization trial was larger than 5 standard deviations of the fUS activity during the session. If there was a vocal production during playback stimulus, the trial was also excluded. We did not use motion correction algorithms to avoid known spurious correlation issues (*46*).

### Number of playback trials (Figs. 1E-I and 2A-F)

After trial exclusion criteria, we obtained a total of 438 (8,130,164,108, 28) phees, 449 (8, 132, 175, 107, 27) trillphees, 428 (8, 134, 150, 108, 28) trills, 453 (8, 133, 177, 108, 27) twitters, 455 (8, 135, 181, 105, 26) alarm calls, and 50 (0, 22, 2, 26, 0) noise stimuli. Each number in the parenthesis indicate the number of playback stimuli for each subject.

### Number of vocal production (Fig. 3B-J) and audio-vocal (Fig. 4A-B) trials

Only three out of five subjects vocalized spontaneously during the experimental sessions. All three subjects produced phee, trill and alarm calls, but not all subjects produced trillphee and twitter. Therefore, we considered phee, trill and alarm calls for vocal production trials. The numbers of calls for each type of call and each subject (indicated in the parenthesis) were the following: spontaneous call phee 173 (23, 35, 115), trill 87 (62, 13, 12), alarm call 61 (23, 14, 24). Response call phee 214 (11, 44, 159), trill 78 (47,1,30), alarm call 85 (25, 18, 42). Sequence call phee 483 (3, 21, 459), trill 81 (21, 0, 60), alarm call 169 (11, 55, 103). For audio-vocal interaction trials, we combined phee and trill as a single group of contact calls.

### Registration of fUS images to a reference image for each subject

We chose a reference session for each subject and calculated the mean intensity at the logarithmic scale, normalized, and converted it to an unsigned 8-bit image. The obtained image was used as the registration template for that subject. The non-brain region above the sagittal sinus was masked out to avoid false signals. We then used the averaged image of each session to align to the reference by Elastix (*47*) (parameter settings see files in ‘registrationParametersFUSi’). We then applied *transformix* to each frame in a session using the *elastix* transformation parameters calculated from the registration step.

### Registration of fUS images of all subjects to a reference subject

To have all the fUS results in a single reference image, we registered all subjects’ images to a single subject. We performed this step after the image registration within each subject. We use the same method as described in “Registration of fUS images to reference images for each subject.”

### Registration of fUS reference to the brain atlas

We carried out a 3-step registration to align the fUS results to a marmoset brain atlas. In the first step, we register the autofluorescent channel of the volumetric light-sheet images of cleared marmoset brains to an MRI atlas (*48*). In the second step, we aligned the midsagittal slice of the co-registered vessel signal channel to the fUS reference using the ImageJ plugin BUnwarpJ (cite “Consistent and Elastic Registration of Histological Sections using Vector-Spline Regularization”). The atlas segmentation of the midsagittal section was thus aligned with the fUS images from the same subject. Available MRI atlas does not have a full annotation of subcortical areas; therefore, we added a third step in which we aligned another marmoset brain atlas (*49*) to the reference brain to complete the annotation. Image registration was done using Elastix. All procedures described bellow were done using the registered fUS images.

### Preprocessing of fUS signal

For each frame, a spatial smoothing using Gaussian kernel with 5 voxel-width and 2 standard deviation was applied (fspecial in MATLAB). For each session, we performed a PCA using all voxels. The first component was related to activities in the large blood vessels; therefore, we excluded the first PCA and reconstructed the whole fUS signal.

### Parcellation of the fUS signal

To describe the brain’s mesoscale activity, we clustered voxels with CBV dynamics that were similar to each other during playback and vocal production separately. To do so, we initially calculated the correlation matrix between each brain voxel. For playback trials, we correlated the CBV activities of each voxel for each stimuli starting from 10s before the stimuli onset up to 30s after the stimuli onset. For vocal production trials, we correlated the CBV activities of each voxel for each stimuli starting from 40s before the stimuli onset up to 40s after the stimuli onset. To obtain the distance matrix between voxels CBV activities, we calculated the FDR corrected p-value for the corresponding correlation. We then calculated the average distance matrix for each type of playback call for each subject. Finally, we averaged over all subjects and types of calls to obtain the overall distance matrix. With this procedure, we make the relevance of each condition and subject on the overall distance matrix balanced. We then used the average distance matrix as input to a spectral clustering algorithm with 100 clusters. Clusters that showed edge artifacts were excluded. The resulting parcellations are shown in figs. S7A-B. The number of parcellation clusters was chosen to approximately match the number of areas annotated for the brain section.

### Calculating the parcellated and mean CBV activity

For each subject and each trial, we calculated the average of CBV activities of the voxels in each parcellation cluster. To obtain the parcellated CBV activity we smoothed the averaged CBV activity using a cubic spline (csap in MATLAB). To calculate the mean CBV activity, we initially averaged the parcellated CBV activities for all trials of interest of each subject and then averaged the CBV activities of all subjects. In this way, we avoided that a single subject biased the mean CBV activity.

### Calculating the significance of the mean CBV (Figs. 1E-I and 3B-F)

For the playback trials, we used the parcellated CBV activity between 10s to 0.5s before the stimuli onset as the baseline. For the vocal production trials, we used the parcellated CBV activity between 30s to 20s before the block stimuli onset as the baseline. This time interval reduced the possibility that the baseline was influenced by CBV activity related to the initiation of vocalization. To calculate the significance of the mean CBV activity for each trial, parcellation cluster, and subject, we calculated the maximum CBV activity during baseline. We then constructed the symmetric 99.95% confidence interval for the maximum baseline values of all trials. We considered that the mean CBV activity for all call trials (phee, trillphee, trill, twitter, alarm call) during playback (0s to 18s after the stimuli onset) was significant if the value was above or below the 99.95% confidence interval (*i*.*e*., p = 0.05 after Bonferroni correction for the 100 parcellated brain areas). We used the Bonferroni correction to guarantee family wise-error rate, so that only the most significant areas were included although there is a risk of missing some weakly activated areas. For Figs. 1E and 3B, we plotted the activity of parcellation cluster which the overlap was largest with the corresponding annotated brain region.

### Calculating the significance of difference between mean CBV (Figs. 2A-E, 3H-I, and 4A-B)

To measure the difference in the dynamics between mean CBV responses for different types of stimuli, we first calculated the mean CBV activity for each type of playback stimuli (for playback trials) and calls (for vocalization and audio-vocal trials). We then calculated the Euclidean distance between the mean CBV response for each call type for the interval 0 s to 30 s with respect to playback onset (for playback trials, Fig. 2), −12 s to 30 s with respect to call onset (for vocalization trials, Fig. 3), and 0 s to 30s with respect to call onset (for audio-vocal trials, Fig. 4). Finally, we multiplied the Euclidean distance with the sign of the difference between mean CBV responses for each parcellation cluster. To calculate the significance of the signed Euclidean distance, we resampled the trials 2000 times and calculated the signed Euclidean distance with a randomized sign to construct the bootstrap statistical test. The randomization of the sign allowed us to construct the distribution for null hypothesis of mean signed Euclidean distance equal to zero. This procedure allowed us to preserve the spatial correlation between different CBV activities, which is ignored in a voxel-wise analysis. We adjusted the p-values using Benjamini-Hochberg FDR correction for the number of parcellation clusters. We used this FDR correction to guarantee high power for the statistical test.

### Hierarchical clustering between CBV activities (Figs. 2F and 3J)

To measure the distance between the CBV responses of the entire medial brain region to different playback stimuli and vocalizations, we calculated the Euclidean distance between call types for each parcellation cluster as described above and averaged among all parcellation clusters. Hence, we generated a distance matrix between the CBV responses for different playback stimuli. We then clustered the brain responses to different call types using the hierarchical clustering algorithm (dendogram in MATLAB).

### Acoustic analysis (fig. S3A-B)

After detecting the onset and offset of the call syllable, a custom-made MATLAB routine calculated the duration, dominant frequency, amplitude modulation (AM) frequency, and Wiener entropy of each syllable (*29*). The duration of a syllable is the difference between the offset and onset of a call. To calculate the dominant frequency of a call, we first calculated the spectrogram and obtained the frequencies at which the spectrogram had maximum power for each time point. The dominant frequency of a syllable was calculated as the maximum of those frequencies. The spectrogram was calculated using an FFT window of 1024 points, Hanning window, with 50% overlap. The AM frequency was calculated in the following way. First, the signal was bandpass filtered between 6 to 10 kHz and then a Hilbert transform was applied. The absolute value of the resulting signal gives us the amplitude envelope of the modulated signal. The 6-10 kHz frequency range was found to give accurate values for the syllable envelope. Finally, the AM frequency was calculated as the dominant frequency of the amplitude envelope. The Wiener entropy is the logarithm of the ratio between the geometric and arithmetic means of the power spectrum values across different frequencies. The Wiener entropy represents how broadband the power spectrum of a signal is. The closer the signal is to white noise, the higher the value of Wiener entropy will be. We reduced the dimensionality by applying PCA to duration, dominant frequency, AM frequency, and Wiener entropy for each call and projecting the values to the first two principal component axis. To calculate the hierarchical clustering, we calculated the mean duration, dominant frequency, amplitude modulation (AM) frequency, and Wiener entropy for each call type. Then we computed the correlation matrix between the call types. We used the correlation matrix as the distance matrix to cluster the call types.

### Correlation matrix between SBN, ACC and motor cortical areas (Fig. 3G, figs. S2, and S5)

To calculate the correlation between the CBV activities of anterior cingulate cortex (Brodmann area 25), anterior hypothalamus, ventromedial hypothalamus, pre-optic area, extended amygdala, lateral septum, periaqueductal gray, limbic thalamus, M1, SMA, and pre-SMA, we first calculated the mean CBV activity for each area during spontaneous vocal production. The registered atlas delineated each area (ROI). Then we correlated the CBV dynamics (from the onset of call to 15s after the call onset) of each ROI to obtain the Pearson correlation matrix.

## Supplementary figures

**fig. S1:**
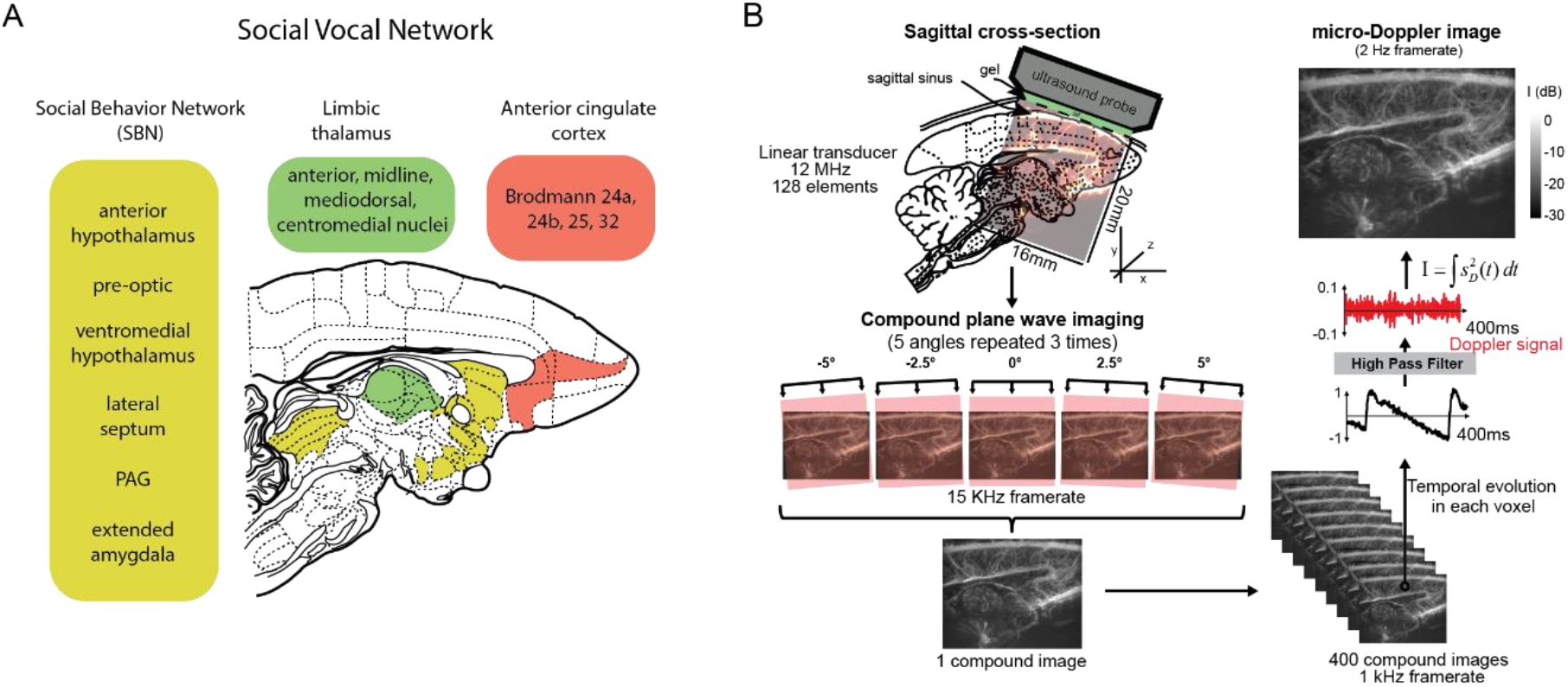
Medial brain region related to social communication. **(A)** Anatomical identification of a social vocal network formed by the social behavior network (SBN), anterior cingulate cortex (ACC) and limbic thalamus (anteromedial, midline, and mediodorsal thalamus). **(B)** Functional ultrasound imaging (fUS) processing of medial brain regions from the ultrasound beam forming to image formation. For details, see (*50*).

**fig. S2:**
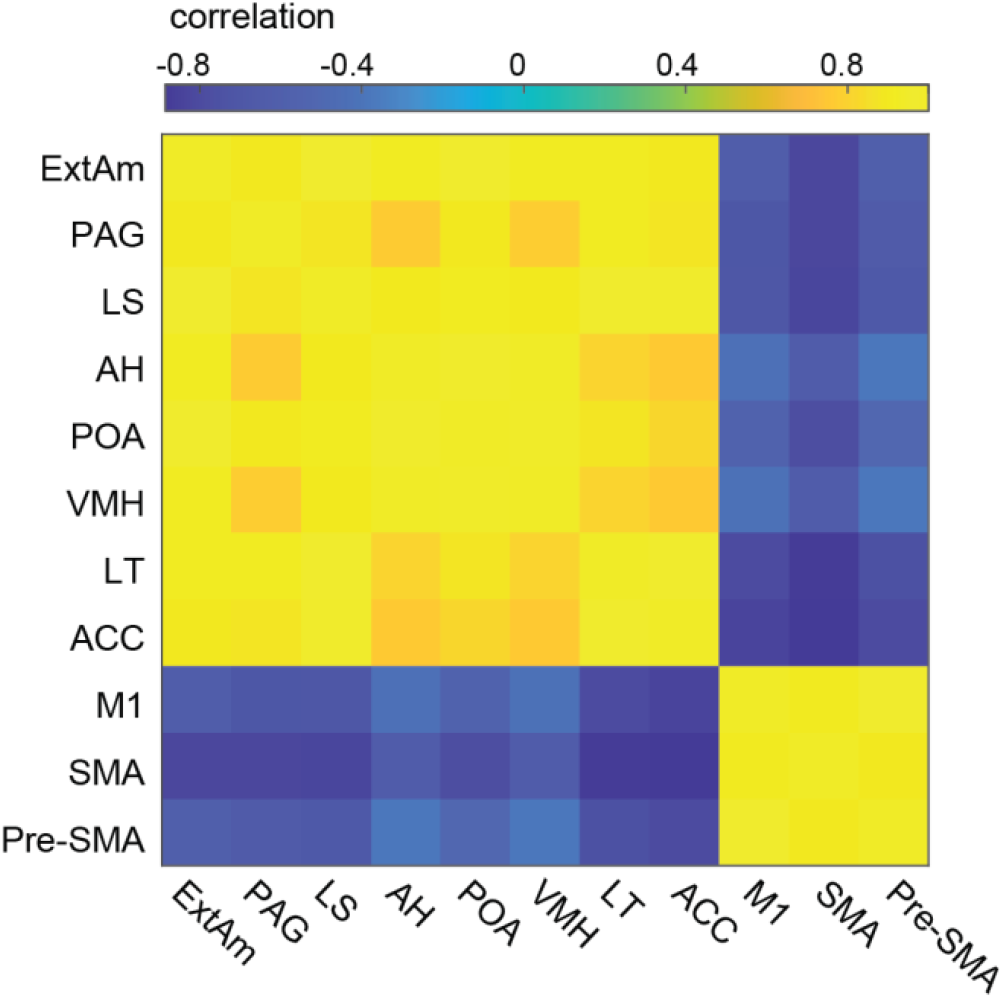
Correlation matrix of the CBV activity in vocal perception. Correlation values between the mean CBV activity (for all stimuli) for each brain area is represented by different colors for the corresponding column and line. All correlations were statistically significant (p<0.05).

**fig. S3:**
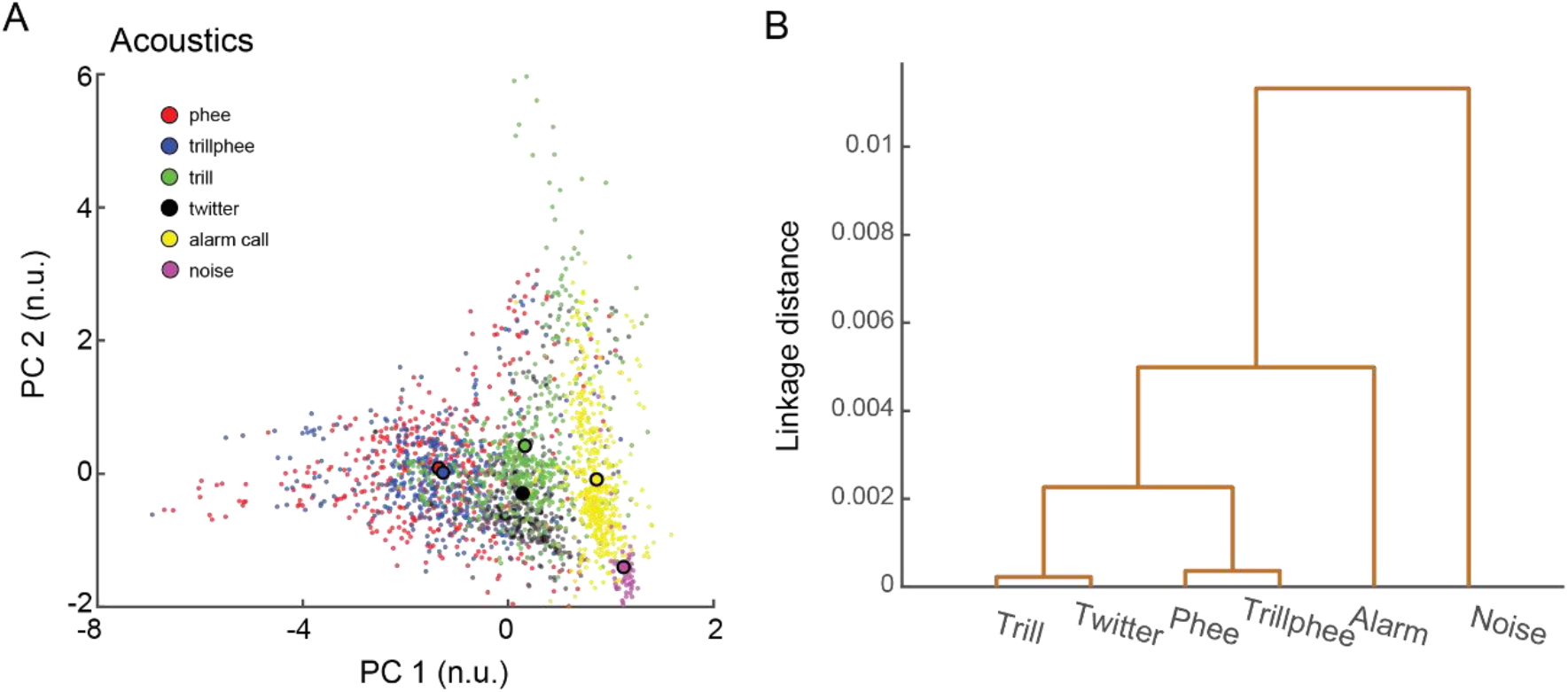
Acoustic characteristics of the playback stimuli and brain response. **(A)** Scatter plot showing the distribution of first two PCA for all playback calls recorded during experiments. PCA of duration, amplitude modulation, dominant frequency, and entropy. **(B)** Hierarchical clustering of the mean acoustic characteristics for each call type.

**fig. S4:**
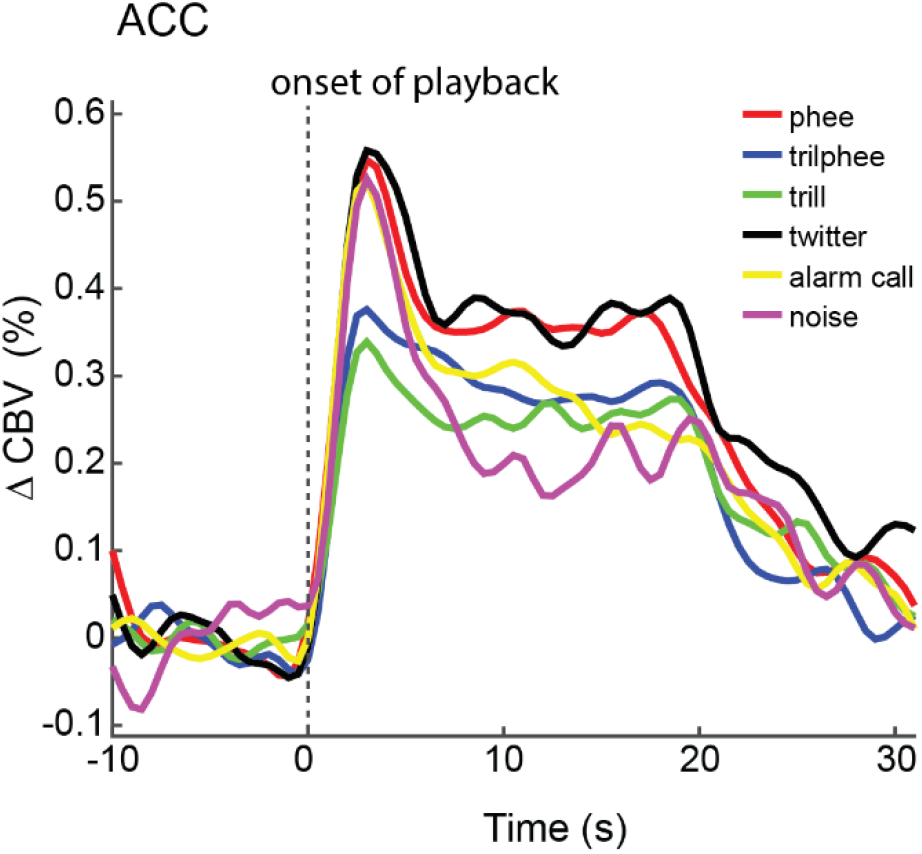
Average CBV response in ACC for each type of call. Observe the difference in the trajectories of the CBV response for different call types.

**fig. S5:**
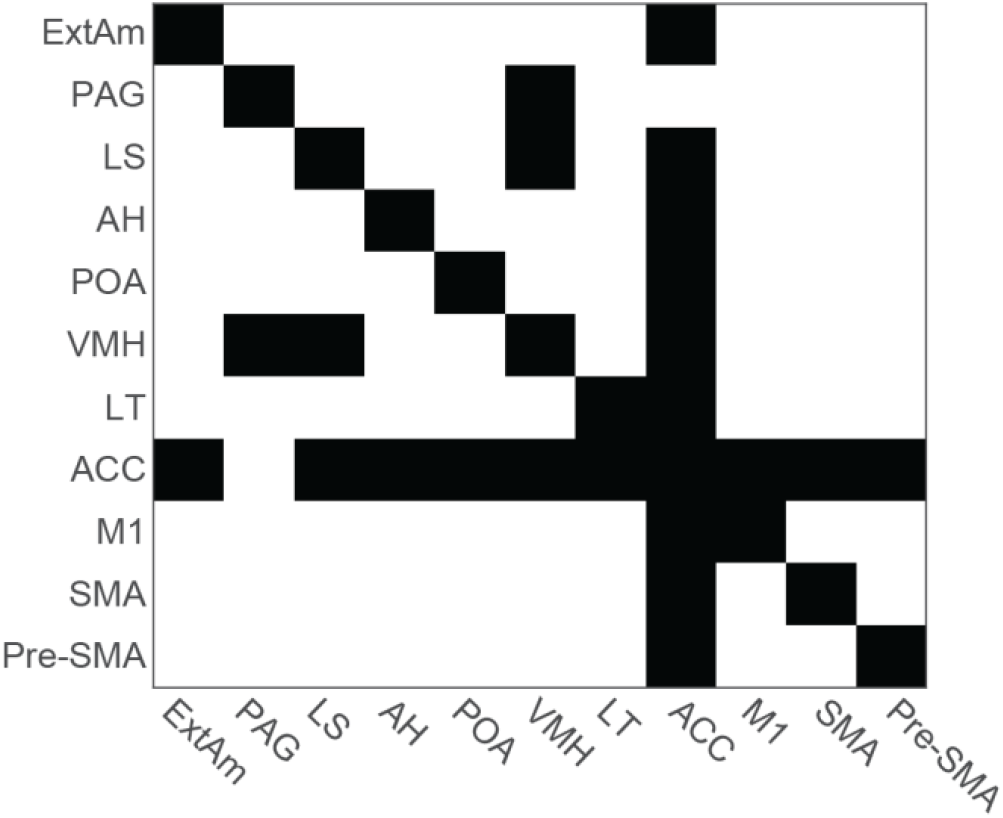
Matrix showing the significant correlations of the CBV activity in vocal production. Significant (p<0.05) and non-significant correlation values are shown in white and black, respectively. The corresponding correlation values are shown in Fig. 3G.

**fig. S6:**
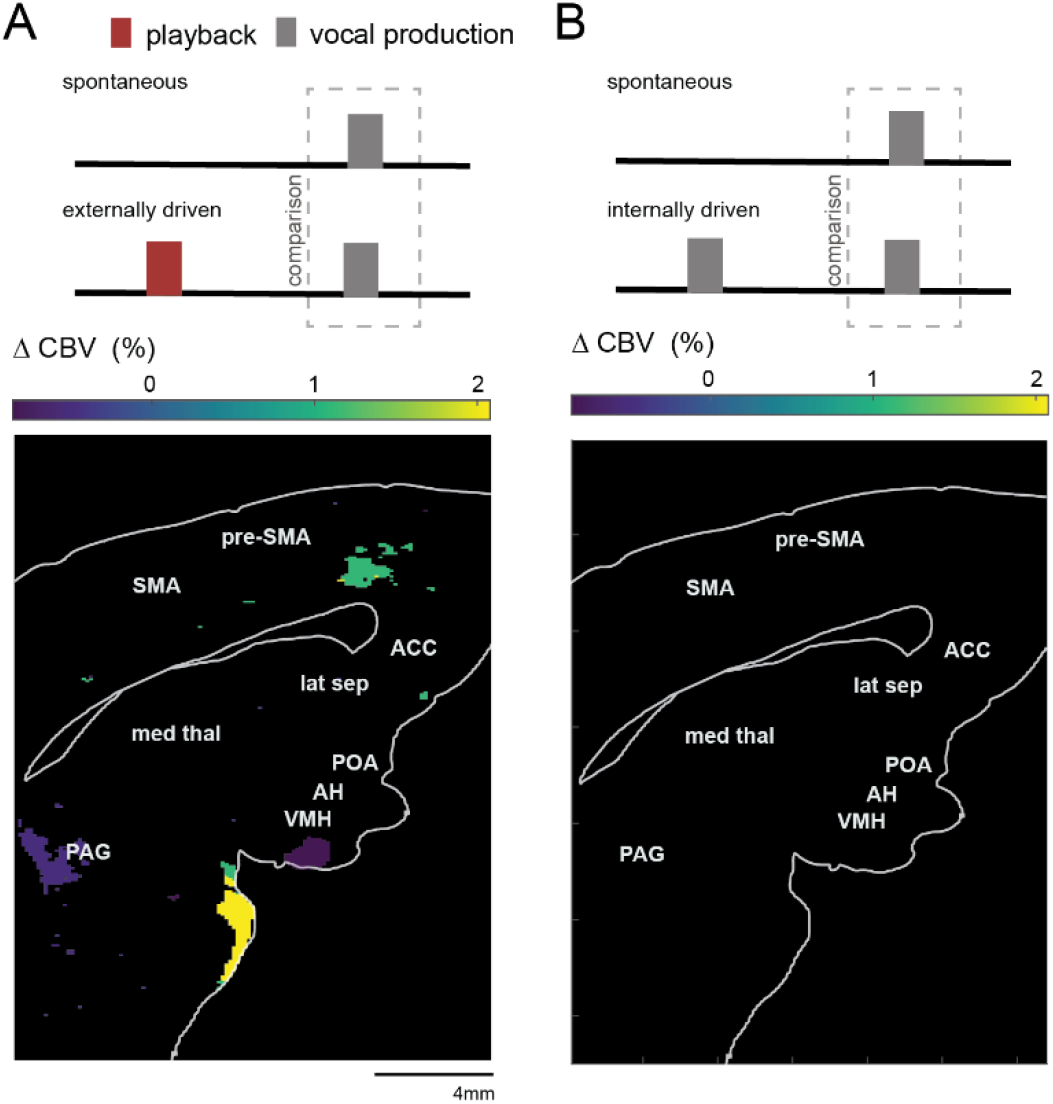
History dependence in the medial brain regions during audio-vocal interaction in alarm calls. **(A)** CBV activity difference between spontaneous and response alarm calls. The brain map shows areas that were significantly different in both conditions (p < 0.05 FDR corrected). Colorbar shows the percentage difference between the CBV activity during production of (externally driven) response and spontaneous alarm calls. Positive values indicate that the activity was stronger for the response to playback calls. **(B)** CBV activity difference between spontaneous and sequence alarm calls. Observe that there was no area with a significant difference.

**fig. S7:**
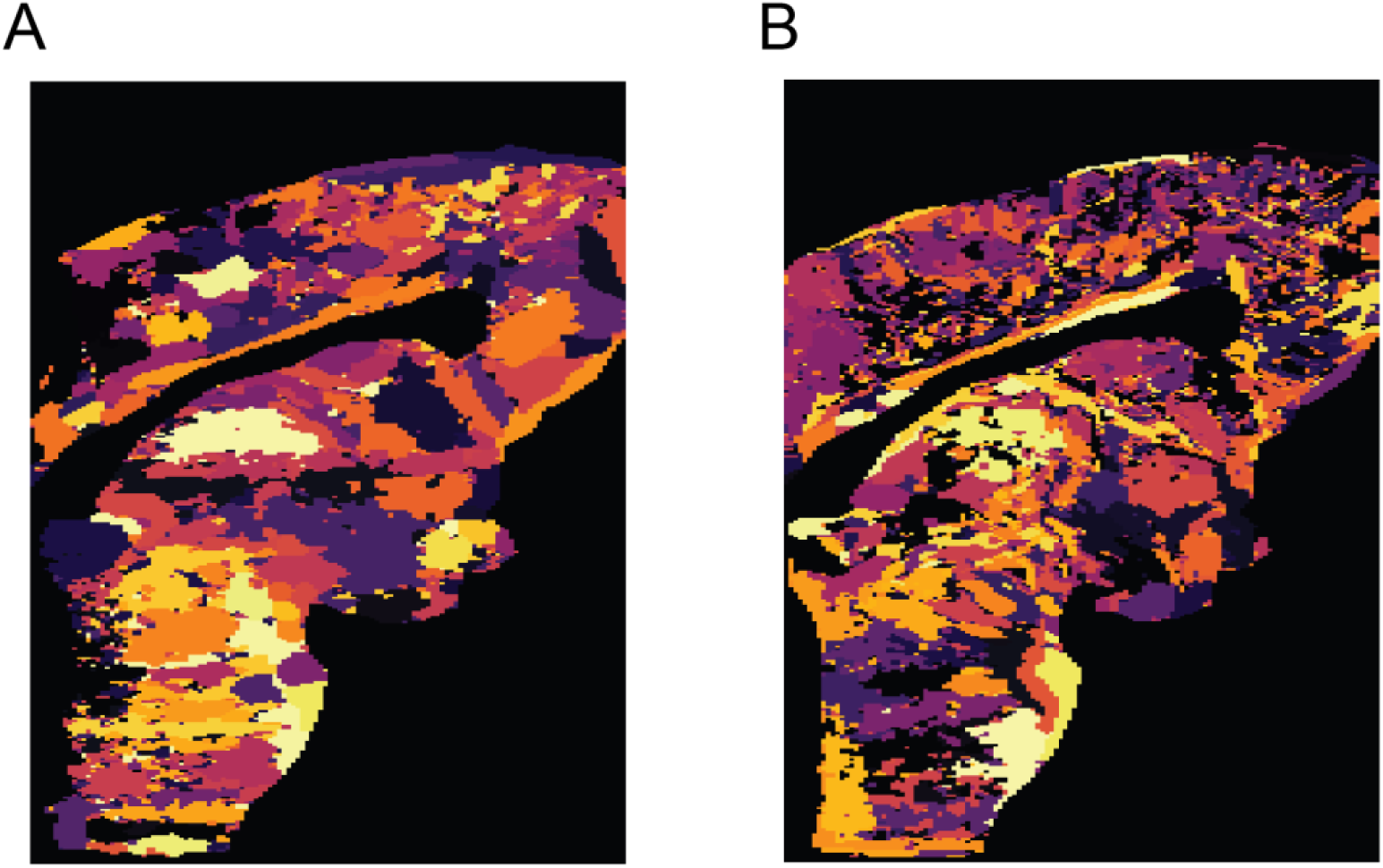
Parcellation of medial brain regions. **(A)** Parcellation result for vocal perception. **(B)** Parcellation result for vocal production.

